# Systematic development of peptide inhibitors targeting the CXCL12/HMGB1 interaction

**DOI:** 10.1101/2019.12.18.878504

**Authors:** Jacopo Sgrignani, Valentina Cecchinato, Enrico M.A. Fassi, Gianluca D’Agostino, Maura Garofalo, Gabriela Danelon, Giovanni Grazioso, Mariagrazia Uguccioni, Andrea Cavalli

## Abstract

During inflammatory reactions, the production and release of chemotactic factors guide the recruitment of selective leukocyte subpopulations. HMGB1 and the chemokine CXCL12, both released in the microenvironment, form a heterocomplex, which exclusively acts on the chemokine receptor CXCR4, enhancing cell migration and, in some pathological conditions such as Rheumatoid Arthritis, exacerbating the immune response. An excessive cell influx at the inflammatory site can be diminished by disrupting the heterocomplex.

Here, we report the computationally driven identification of a novel peptide (HBP08), which binds HMGB1 with the highest affinity reported so far (K_d_ of 0.8 ± 0.1 μM), able to selectively inhibit the activity of the CXCL12/HMGB1 heterocomplex.

The identification of this peptide represents an important step towards the development of innovative pharmacological tools for the treatment of severe chronic inflammatory conditions characterized by an uncontrolled immune response.

## Introduction

Chemokines are key regulators of leukocyte migration and play fundamental roles both in physiological and pathological immune responses.^1^ Chemokine receptors differentially expressed by all leukocytes and many non-hematopoietic cells, including cancer cells, constitute the largest branch of the γ subfamily of rhodopsin-like G protein-coupled receptors (GPCR). In modern pharmacology, this receptor superfamily represents the most successful target of small molecule inhibitors for the treatment of a variety of human diseases.^2^ In the last 25/30 years, an impressive amount of preclinical and clinical evidence has progressively validated the role of chemokines and their receptors in immune-mediated diseases.^3, 4^

Furthermore, in the last decade, several studies have pointed out how the activity of chemokines on cell migration can be modulated by their binding to other chemokines or proteins released in inflammation.^5, 6^ In particular, our group has shown that High Mobility Group Box 1 (HMGB1), an alarmin released under stress conditions, forms a heterocomplex with the chemokine CXCL12, favoring cell migration via the activation of the chemokine receptor CXCR4 in the presence of low concentration of CXCL12, which normally is insufficient to trigger a cellular response.^7^ Moreover, we have demonstrated that synergism between CXCL12 and HMGB1 sustains inflammation in Rheumatoid Arthritis (RA).^8^

These observations suggest that the identifications of molecules able to suppress the interaction between chemokines and their modulators could lead to the discovery of effective inhibitors able to promote resolution of inflammation.

HMGB1 is a highly conserved nuclear protein expressed in bacteria, yeast, plants and in all vertebrate cells. Structurally, it is composed by two homologous, but not identical domains, BoxA and BoxB, and a negatively charged C-terminal tail (Figure 1).^9^ In addition to its nuclear function, HMGB1 is released under inflammatory conditions or by necrotic cells, and acts as a damage-associated molecular pattern molecule (DAMP).^10, 11^

**Figure 1.**
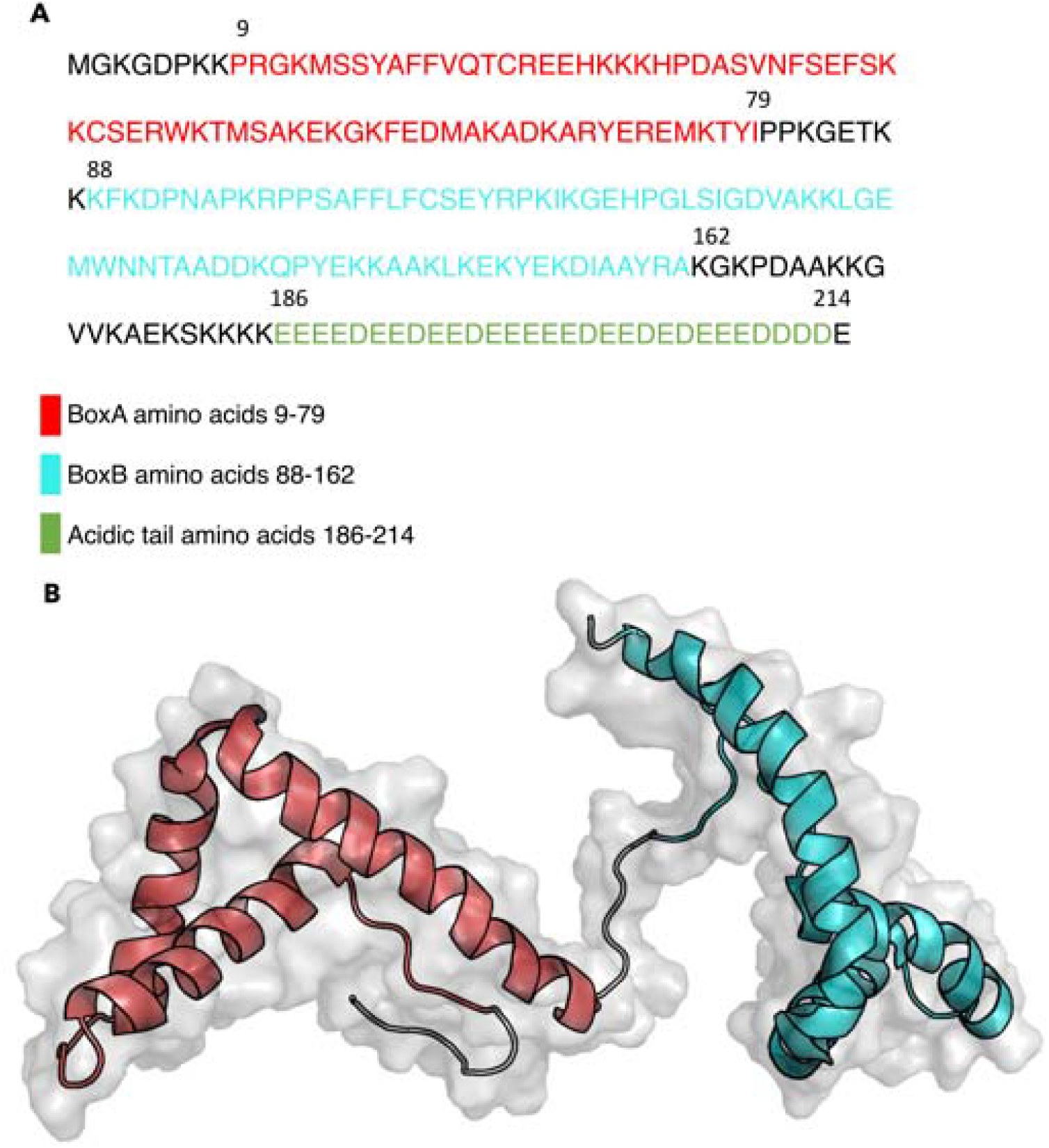
(**A**) Amino acid sequence of HMGB1. Residues constituting the two boxes are shown in red (BoxA) and cyan (BoxB), while the acidic tail is shown in green. (**B**) Ribbon representation of the two boxes of HMGB1 structure in solution (fragment 2-174, PDB code 2YRQ).

In the extracellular space, HMGB1 can be present in different redox states, depending on the presence of an intramolecular disulfide bond between two cysteines at position 23 and 45.^12^ Reduced HMGB1, once released in the extracellular space, can form a heterocomplex with CXCL12 and synergistically promote, via CXCR4, the recruitment of leukocytes to inflammatory sites.^7, 8, 13^ Moreover reduced HMGB1 can bind to the receptor for advanced glycation endproducts (RAGE) to induce CXCL12 secretion and authophagy.^14^ Once oxidized, by reactive oxidative species present in the extracellular space, HMGB1 binds to the Toll-like Receptor 4 (TLR4) leading to activation of the nuclear factor kappa-B (NF-kB) and transcription of cytokines, and chemokines.^12, 15^

To date, despite the importance of this target, only few inhibitors of the CXCL12/HMGB1 interaction, or of the HMGB1 functions have been identified.^16, 17, 18, 19^ Currently, glycyrrhizin is the most potent and the best structurally characterized inhibitor of the CXCL12/HMGB1 heterocomplex, but has a low affinity for HMGB1 (K_d_ ~ 150 μM), and it lacks of specificity.^7, 16, 19^

Peptides are receiving increasing attention due to their ability in targeting large surfaces as those involved in protein-protein interactions (PPI), and to promising pre-clinical and clinical results.^20,22^ Recent efforts have been put into the development of innovative strategies to overcome their intrinsic limitations such as low bioavailability and poor metabolic stability.^21, 22, 23^ It is estimated that more than 400 peptides are in clinical development, and 60 are already available for therapeutic use in different countries.^24, 25^

Here, we report the computationally driven identification of a novel high affinity nonapeptide able to inhibit the formation of the CXCL12/HMGB1 heterocomplex and to abolish the synergistic effect on cell migration in CXCR4 transfected cells and in human monocytes, without affecting the ability of HMGB1 to trigger TLR4. The peptide is the strongest HMGB1 binder reported so far, with an affinity K_d_ of 0.8 µM.

## Results

### Design of a peptide inhibitor of the CXCL12/HMGB1 interaction

Taking advantage of the known interaction between glycyrrhizin and HMGB1,^16^ we developed a computational pipeline to identify novel and selective peptide inhibitors of the CXCL12/HMGB1 interaction (Figure 2A).

**Figure 2.**
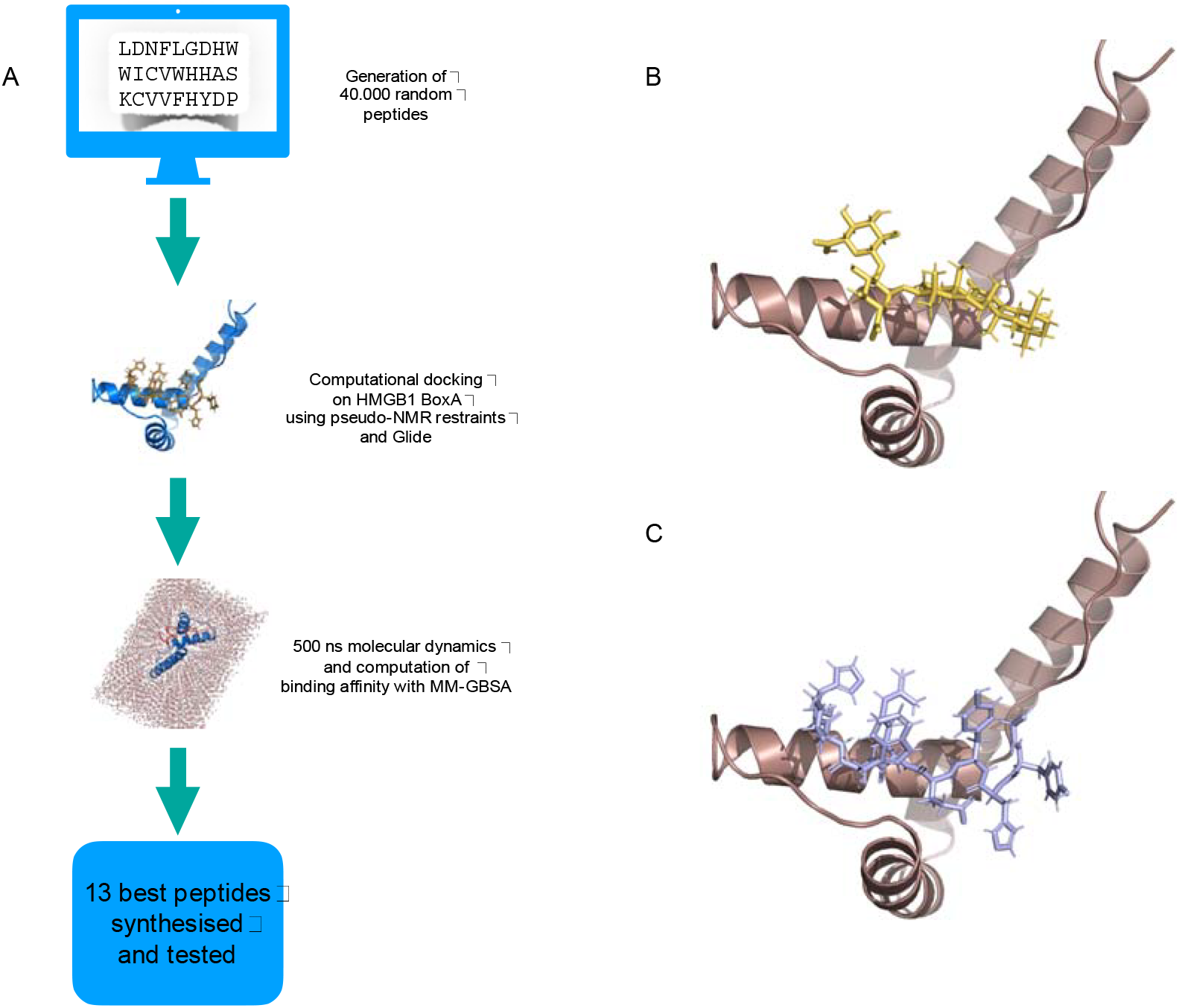
(**A**) Workflow diagram of the computational pipeline used for the identification of the binding peptides. Peptides with a randomly generated sequence were first docked using pseudo-NMR restraints and then re-docked with Glide. Finally, peptides were ranked according to their binding free energy (∆G) computed using MMGBSA with explicit water simulations of 500ns. (**B**) Model of the glycyrrhizin-BoxA complex used to define the peptide binding site. (**C**) Model of the complex of one of the identified peptides (HBP08) with BoxA obtained after the first docking.

We generated a model of the glycyrrhizin/BoxA complex consistent with the results of previously reported NMR chemical shift perturbation studies (Figure 2B).^16^

To maximize the heterogeneity of the peptides considered in our screening, we generated a library of 40.000 nonapeptides with a randomly selected sequence. All peptides were docked in the glycyrrhizin binding site and ranked according to the binding energy of the corresponding peptide/HMGB1 complex (Figure 2C, See Materials and Methods). Finally, aiming to reduce the number of potential false positives, the best 100 ranking peptides were re-docked to BoxA, with the program Glide,^26^ without constraining the algorithm to explore only the glycyrrhizin binding site.

The peptides resulting after these calculations were visually inspected and only the best GSCORE (a scoring function aimed to estimate binding affinity) pose of 57 peptides with a glycyrrhizin-like binding mode were retained for further analysis (Supplementary Table 1).

Several studies have shown that approximated free energy methods like MM-GBSA,^27, 28^ especially when coupled with long MD simulations, can be a valuable help in selection of active molecules in virtual screening investigations.^29, 30^ Therefore, a 500 ns long MD simulation was performed for each of the 57 peptides obtained from docking calculations. Those detaching from the BoxA binding pocket during the simulations (14 out of 57) were considered unstable and not further analyzed in MM-GBSA calculations (Supplementary Table 1).

Based on the MM-GBSA score, 13 different peptides were selected to be tested *in vitro* (Table 1, and Supplementary Figure 1).

**Table 1.**
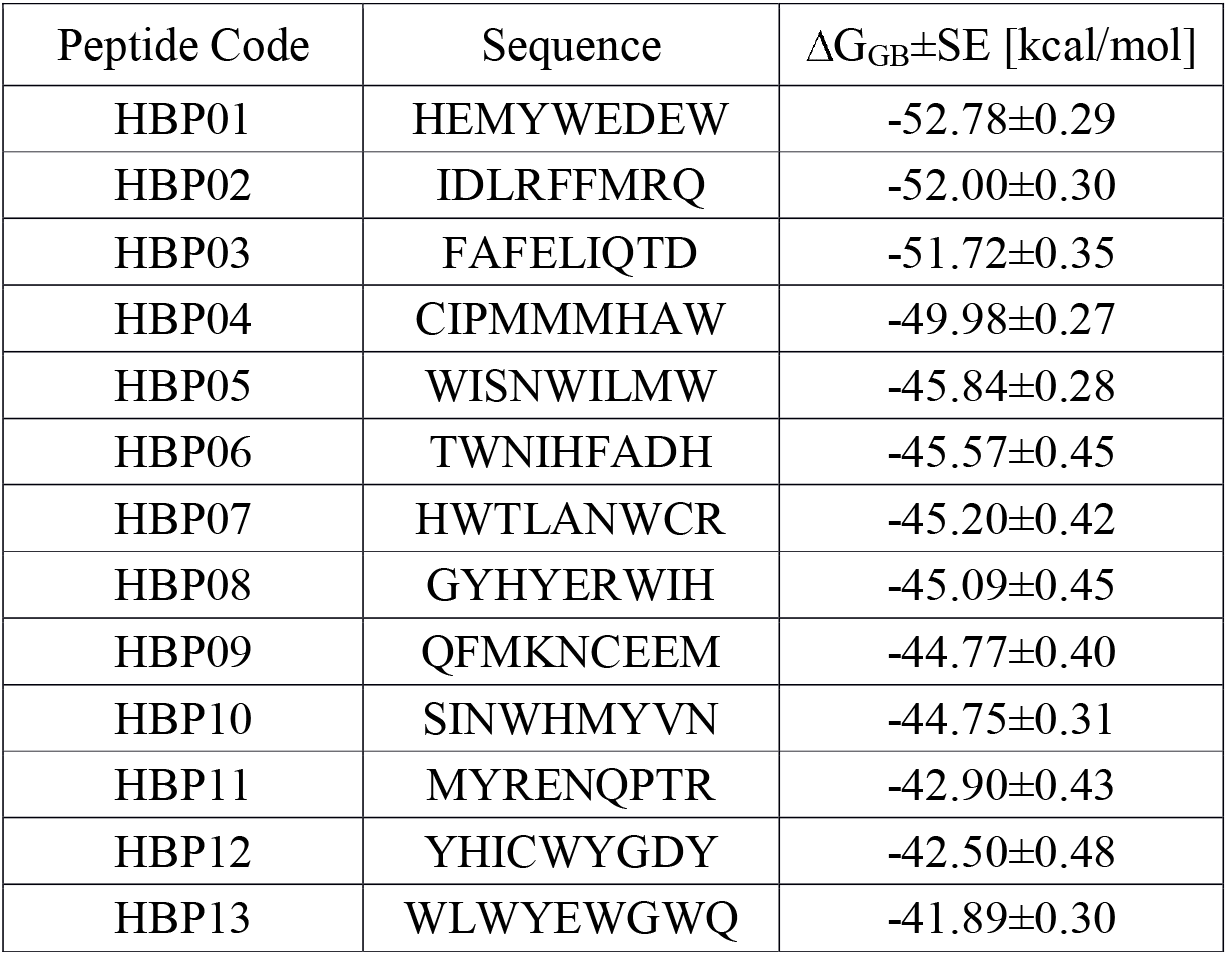
List of binding peptides ranked according to their theoretical binging free energy ∆G.

### In vitro assessment of the identified peptides

The 13 identified peptides were tested in *in vitro* chemotaxis assay, to evaluate their efficacy as inhibitors of the CXCL12/HMGB1-induced migration, on a murine cell line expressing the human CXCR4. Our experiments showed that 4, out of 13 peptides tested, efficaciously inhibited the enhanced migration induced by the CXCL12/HMGB1 heterocomplex (Figure 3A). Of note, the inhibition observed using 100 µM of HBP05, HBP07, HBP08, or HBP12 was similar or better than the one observed using glycyrrhizin at 200 µM (Figure 3A). Further experiments performed with CXCL12 alone, showed that HBP07 and HBP08 do not affect CXCL12-induced cell migration, while HBP05 and HBP12 inhibit the migration induced by the chemokine alone (Figure 3B), and therefore were not used for further experiments. HBP07 and HBP08 were then tested on primary human monocytes. Only the HBP08 significantly blocked the activity of the heterocomplex (Figure 3C), without altering the migration induced by CXCL12 alone (Figure 3D), and exhibited no toxicity on both cell types (data not shown).

**Figure 3.**
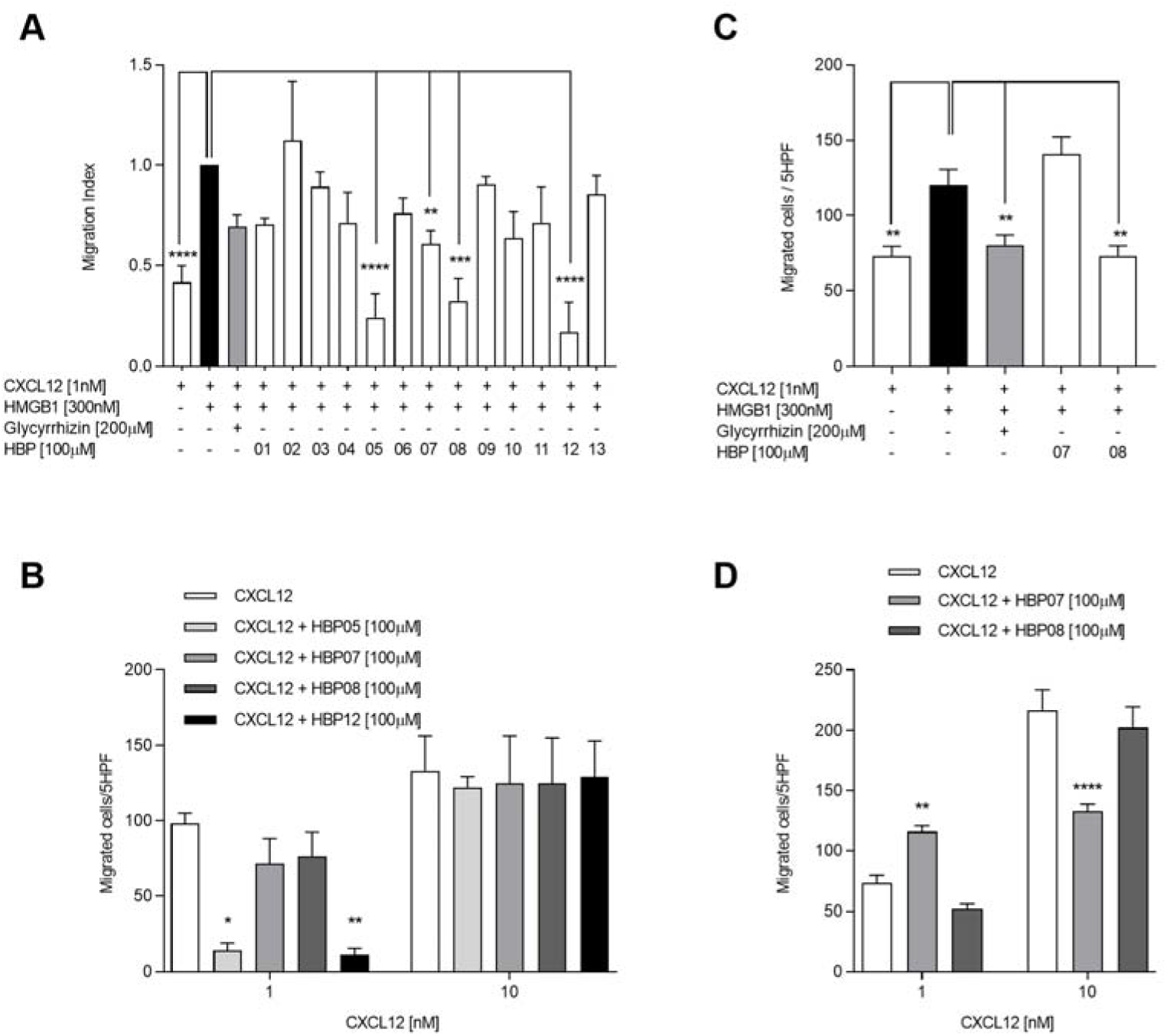
*In vitro* activity of the identified peptides. (**A**) Inhibition of cell migration in response to the CXCL12/HMGB1 heterocomplex was assessed on 300-19 Pre-B cells transfected with human CXCR4 using the identified peptides or glycyrrhizin. Migration index was calculated as the ratio between the number of cells migrated in response to the heterocomplex in the presence or absence of the peptides. (**B**) Migration induced on 300-19 Pre-B cells transfected with CXCR4 by CXCL12 alone in the presence or absence of the peptides identified in (A) as inhibitors of the migration induced by the heterocomplex. (**C**) Inhibition of cell migration in response to the CXCL12/HMGB1 heterocomplex was assessed on human monocytes using HBP07, HBP08, or glycyrrhizin. (**D**) Migration induced on monocytes by CXCL12 alone in the presence or absence of HBP07, HBP08. (A-D) Migrated cells were counted in 5 high-power fields (HPF), and data are shown as mean±SEM of at least three independent experiments performed. *p<0.05; **p<0.01; ***p<0.001; ****p<0.0001 by one-way ANOVA followed by Dunnett’s multicomparisons test (A, C), or two-way ANOVA followed by Tukey’s multicomparisons test (B, D).

### Selective activity of the HBP08 peptide

In the extracellular space oxidized HMGB1, through the binding to TLR4, activates the NF-kB pathway and induces the transcription of several pro-inflammatory cytokines.^12, 15^ In order to determine whether HBP08 was a selective inhibitor of the activity of the CXCL12/HMGB1 heterocomplex or could also prevent the binding of HMGB1 to its receptor TLR4, we performed a cytokine release assay on monocytes treated with HMGB1 alone, or in the presence of HBP08. Monocytes stimulation with HMGB1 induced a significant release of IL-6 and TNF, which could be blocked by treatment with a neutralizing antibody against TLR4 (Figure 4A, B). HBP08 did not induce IL-6 or TNF release and did not block the HMGB1-mediated release of these cytokines. These data indicate that HBP08 selectively inhibits the CXCL12/HMGB1 heterocomplex activity, leaving HMGB1 able to trigger TLR4.

**Figure 4.**
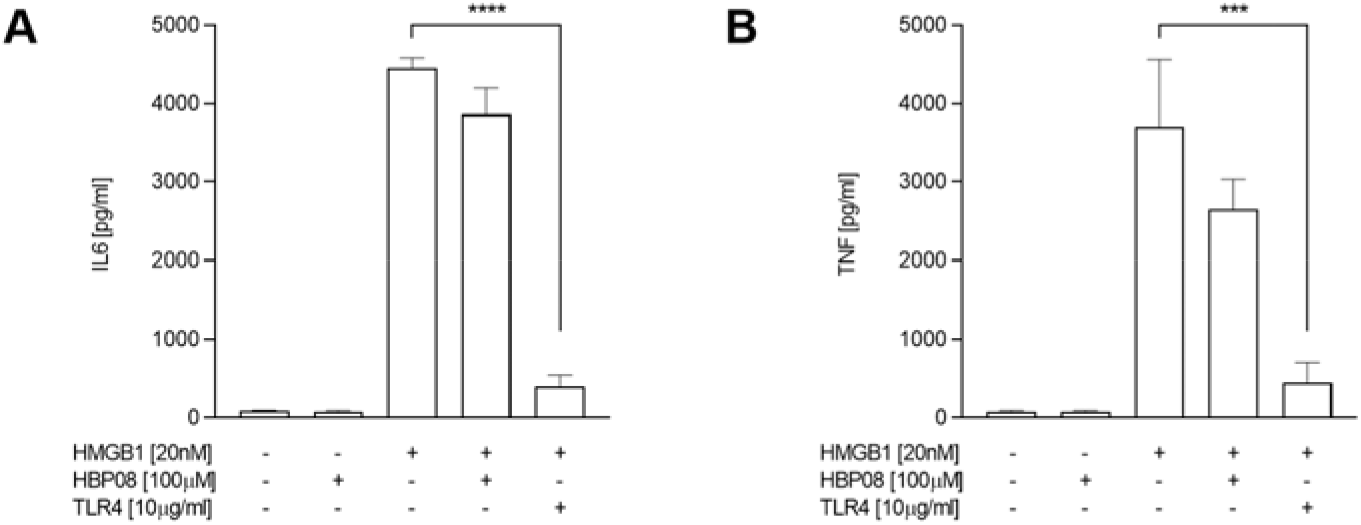
HMGB1-induced release of IL6 and TNF via TLR4 is not inhibited by HBP08. The concentration of IL6 (**A**) and TNF (**B**) in the supernatant of monocytes treated with HMGB1 or LPS in the presence of HBP08 or a neutralizing antibody against TLR4 was measured by CBA. Data are shown as mean±SEM of at least four independent experiments performed. ***p<0.001; ****p<0.0001 by one-way ANOVA followed by Dunnett’s multicomparisons test.

### Characterization of the HMGB1-HBP08 interaction

Microscale thermophoresis (MST) was performed to determine the affinity of HBP08 to HMGB1, resulting in a K_d_ of 0.8±0.1 μM (Figure 5). The affinity of the identified peptide is therefore two orders of magnitude higher than the reported value for glycyrrhizin (K_d_~150 µM).^16^ Overall, these results indicate HBP08 as the first selective and potent peptide inhibitor of the CXCL12/HMGB1 heterocomplex, developed so far.

**Figure 5.**
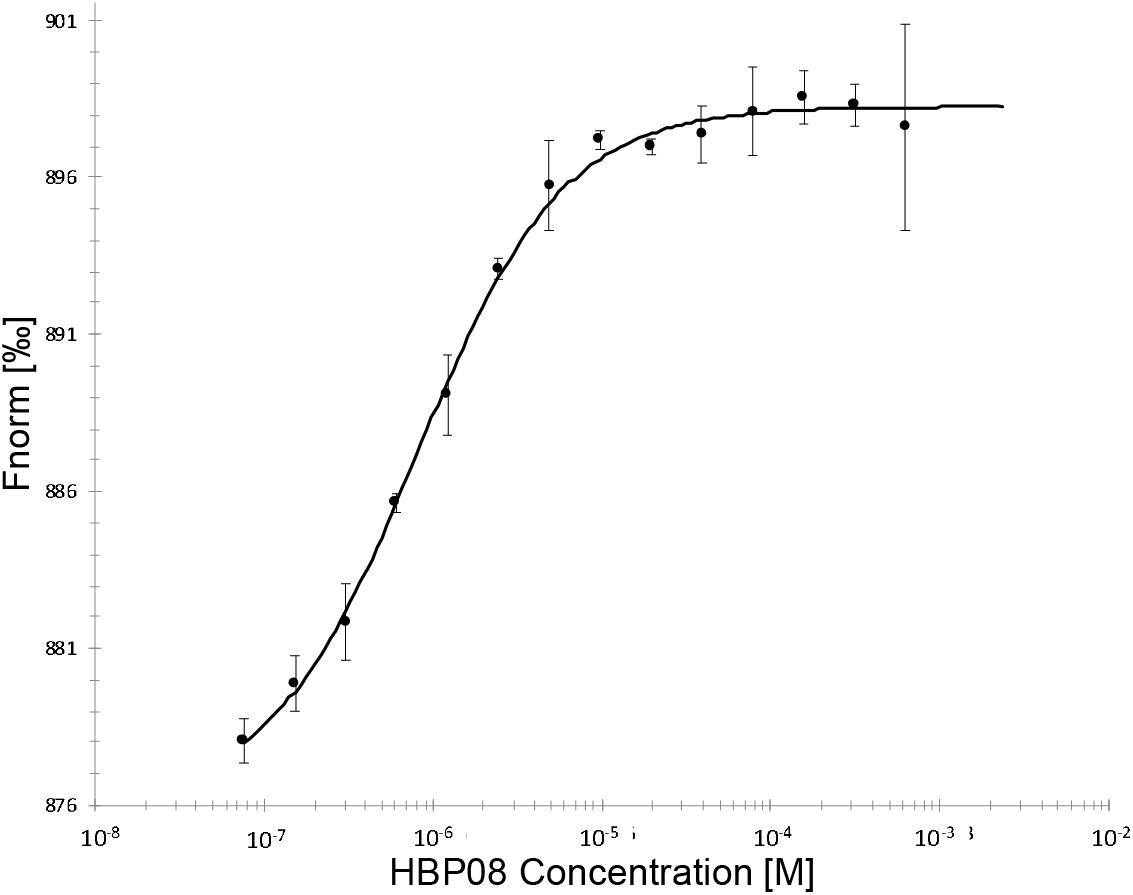
Microscale thermophoresis analysis of the interaction between HBP08 and HMGB1 (K_d_ = 0.8 ± 0.1 μM).

To further characterize the interaction between HBP08 and HMGB1, we performed both experimental and computational alanine scanning of the peptide, comparing the binding affinities of the mutants measured by MST with the change in free energy (∆∆G) estimated by the computational procedure implemented in BioLuminate.^31, 32^ (Table 2).

**Table 2.**
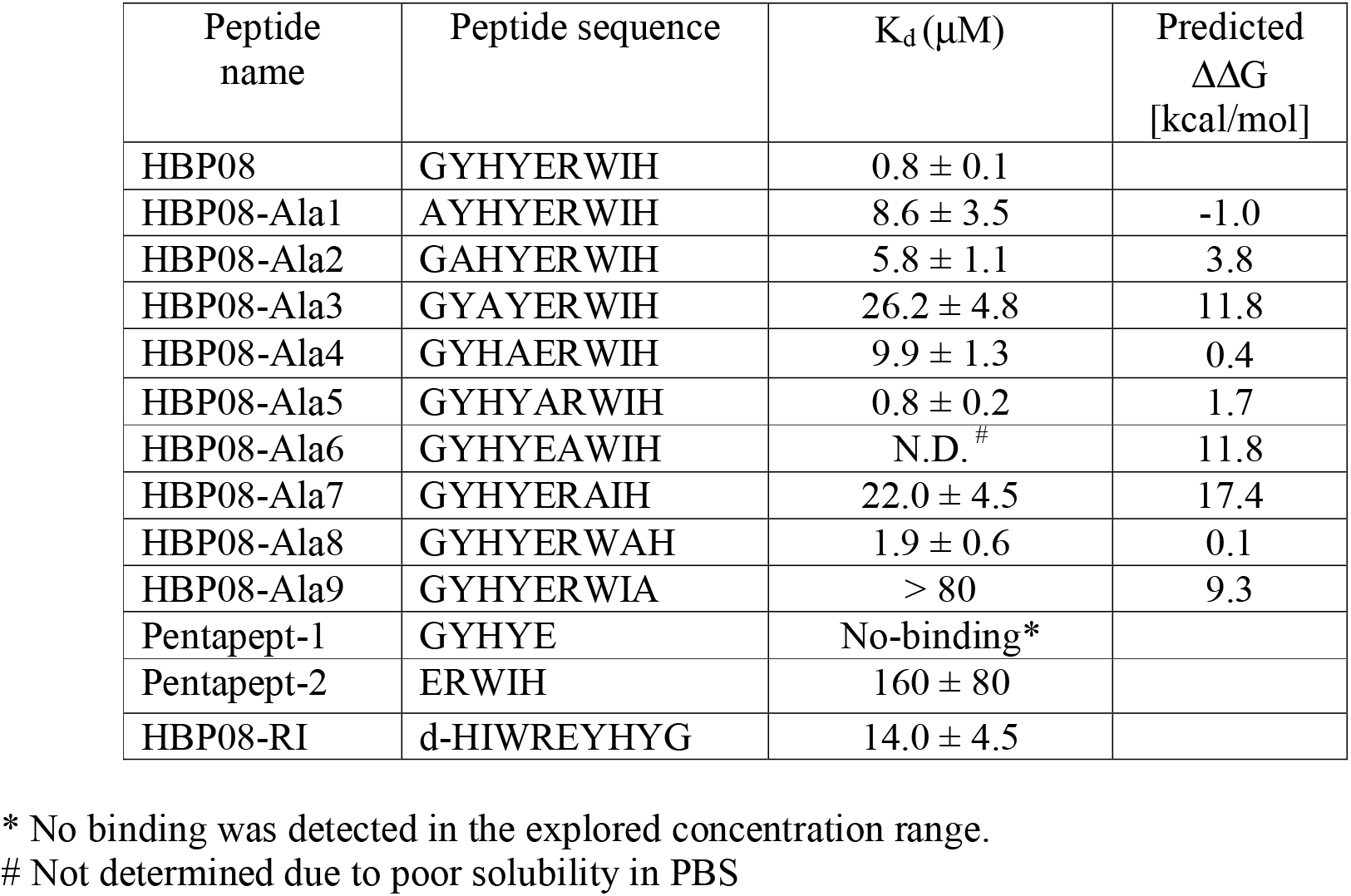
Equilibrium dissociation constant (K_d_) and predicted binding energy (ΔΔG) for the complexes between HMGB1 and the peptide of the first column

The comparison between the experimentally determined K_d_ and the predicted ΔΔG values showed a good agreement. Out of the four mutations with the highest positive binding ∆∆G value predicted (HBP08-Ala3, HBP08-Ala6, HBP08-Ala7 and HBP08-Ala9), three showed a clear decrease in the affinity for HMGB1 measured by MST (HBP08-Ala3, HBP08-Ala7 and HBP08-Ala9). A high positive ∆∆G was estimated also for HBP08-Ala6, in which an arginine residue is mutated in alanine, however this change resulted in a poorly soluble peptide which affinity could be not determined by MST. Concerning the other HBP08 residues, for which smaller binding ∆∆G were predicted, their K_d_ values resulted similar to the value measured for HBP08.

Finally, we also tested the affinity of two peptides formed by the first (pentapept-1) or the last (pentapept-2) five residues of HBP08. Concerning pentapept-1, we did not observe binding in the range of concentration applied to the analysis of the other peptides, while a K_d_ of 160±80 µM was determined for pentapept-2. In summary, both computational and experimental analysis of the binding determinants of HBP08 indicated that the residues at the positions 3, 7 and 9 are the most important for the binding.

To further understand the structural determinants of the individual contributions of these key residues to the binding of the peptide, we analyzed the 3D structure of the HBP08/HMGB1 complex (Figure 6A, B). From this analysis, HBP08-Arg6 and HBP08-Trp7 form h-bond interactions with Asp67, while HBP08-His9 establishes the same type of interaction with Arg24. Differently, HBP08-His3 is placed in a cavity delimited by Ala17, Val20 and Arg24 and its contribution to the binding seems to be mainly due to VdW interactions.

**Figure 6.**
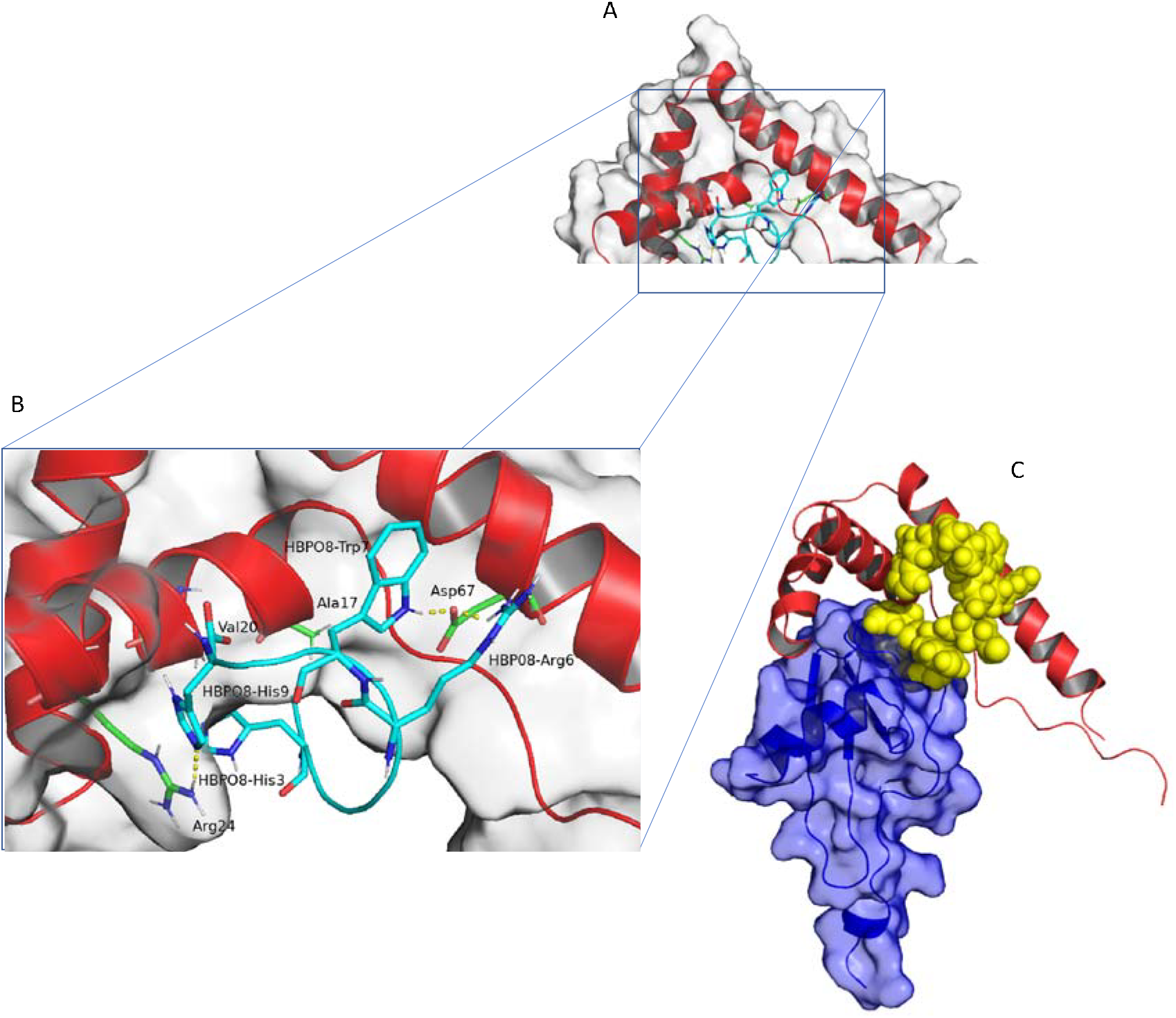
**(A)** Molecular model of the HBP08-BoxA complex. BoxA and HBP08 are represented as red or aquamarine cartoons, respectively. The residues more important for the binding are represented as sticks colored by atom type. (**B**) Focus on critical interactions between HBP08 and BoxA. (**C**) Comparison between the HBP08 binding mode and the structure of CXCL12-BoxA complex obtained by docking.

Finally, we compared the structure of the HBP08/BoxA complex with the one of the CXCL12/BoxA complex, obtained integrating previous NMR investigations^7^ and computational modeling^13^ (Figure 6C). This analysis disclosed that the HBP08 binding site on BoxA is formed by some residues Ala17, Val20, Arg24 and Glu25, important also for the CXCL12 binding.^13^ Therefore, the peptide binding might antagonize by competition the formation of the CXCL12/HMGB1 heterocomplex.

### HBP08 retro-inverso

L-peptides are susceptible to the action of proteolytic enzymes such as peptidases, hindering their application *in vivo*. D-peptides are less prone to the action of peptidases and to the acidic hydrolysis that occurs in the stomach, which increases their oral bioavailability and half-live in the blood circulation. Furthermore D-peptides have a lower immunogenicity.^33^ Taken together, all these features make D-peptides more suitable for drug development.^34^

To exploit the potential of D-peptides, we investigated the binding of a retro-inverso analog of HBP08 (HBP08-RI) made by D-amino acids in reversed order. The results of the binding experiments indicated that HBP08-RI has a good affinity for HMGB1 (K_d_ = 14.0 ± 4.5 µM) and represents, therefore, a good candidate for future drug development studies.

## Discussion

Over the last years, several reports have demonstrated the relevance of the CXCL12/HMGB1 heterocomplex both in physiological and in pathological processes. Recently, the CXCL12/HMGB1 heterocomplex has been shown to be crucial in the perpetuation of the chronic inflammation observed in RA, by fueling the recruitment of immune cells.^8^ Several therapeutic approaches based on the use of biologic and synthetic therapies are currently in use for the treatment of RA, but a portion of patients does not benefit of the available treatments and only the 20-30% of them reach a low disease activity status.^35, 36^ Interestingly, Pitzalis and coworkers have recently pointed out that the composition of the synovial tissue of patients with RA can be related with the response to therapies.^35^

We have recently shown that the CXCL12/HMGB1 heterocomplex is present in the synovial fluids of patients affected by RA, and that its function is maintained in patients with active disease.^8^ Therefore, small molecules or peptides able to hinder the formation of this heterocomplex could be useful as novel personalized therapeutic strategies.

Multiple attempts have been made to identify small molecules able to bind HMGB1.^19^ The majority of inhibitors reported in literature so far show a weak affinity for HMGB1, and are not selectively targeting its synergistic interaction with CXCL12.^16^ Recently, diflunisal has been reported as a specific inhibitor of the CXCL12/HMGB1 heterocomplex activity, without affecting the TLR4 signaling, but its Kd for HMGB1 is only in the mM range.^18^

Peptides are considered a class of molecules particularly suitable to target protein-protein interaction and they are attracting a renewed interest by medicinal chemists.^37^ Therefore, we have applied a computational pipeline to design peptides able to inhibit the formation of the CXCL12/HMGB1 heterocomplex.

Out of the 13 candidates selected with the computational procedure, HBP08 resulted to be able to efficiently inhibit the synergy induced by the heterocomplex on murine cells transfected with the human CXCR4 and on human monocytes. HBP08 and its retro-inverso version bind to HMGB1 with an affinity greater than glycyrrhizin or diflunisal, representing new molecular tools to be exploited for further investigating *in vitro* and *in vivo* the role of the CXCL12/HMGB1 heterocomplex in different inflammatory conditions.

Previous studies of Al-Abed and coworkers^38^ indicated that the TLR4 activation by HMGB1 can be inhibited by both BoxA and an anti-HMGB1 antibody (2G7) that interacts with HMGB1 binding to the region within the residues 53-63 of BoxA.^39^ These results indicated that the same region, far from those we identified for the HBP08 binding, should be responsible of the HMGB1/TLR4 interaction, and in fact we have demonstrated that the developed peptide does not influence the HMGB1 functions related to the TLR4 axis.

The results presented here, demonstrate how the applied computational pipeline allows the fast and efficient design of peptides able to antagonize protein-protein interaction. We propose its application as a novel strategy for the development of powerful inhibitors of protein-protein complex formation.

## Methods

### Glycyrrhizin docking to HMGB1

A model of the HMGB1-glycyrrhizin complex was built by ligand docking, starting from NMR HMGB1 structure available in the protein data bank with the code 2YRQ. All the docking calculation were carried out using Glide (Schrodinger Inc.) in the version 2016-4.^40^ The grid necessary to perform docking was centered in the COG (center of geometry) of the protein and both the enclosing and the bounding box were set bigger than entire protein, to allow a blind-docking, i.e. docking without previous knowledge of a binding site. Standard precision (SP) mode was used to score the resulting ligand-protein complexes.

The twenty poses with the best Glide score were kept for further investigation. Finally, the structure with the best agreement with NMR chemical shifts perturbation (CSP) data by Mollica et al.^16^ was selected as the most likely representative model of the HMGB1-glycyrrhizin complex.

### Computational design of binding peptides

Peptides were designed following a multistep process. First, the model of the BoxA-glycyrrhizin complex was used to define the target binding site for the peptides. To this end, we selected all amino acids from BoxA for which at least a carbon atom was at a distance smaller than 7.5 Å from a glycyrrhizin carbon atom. These gave a list of 17 amino acids, namely: LYS_12, MET_13, SER_14, SER_15, TRY_16, ALA_17, VAL_20, GLU_21, ARG_24, GLU_25, LYS_28, SER_35, VAL_36, ASN_37, PHE_38, PHE_41, SER_42.

Since the size of glycyrrhizin is approximatively equal to the length of a linear 9-residue peptide we proceeded with the generation of 40,000 9-residue peptides with a random sequence. All these peptides were then docked on the BoxA domain using the torsional angular molecular dynamics (TMD) module^41, 42^ of the software package ALMOST.^43^

The docking of the peptides was guided by a set of 17 synthetic NMR-like ambiguous upper-distance restraints^44^ between the Cα atoms, *i*, of the residues of the binding site of BoxA and the Cα atoms, *j*, of the peptide,

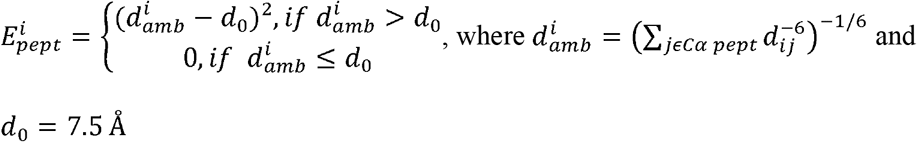

For each peptide, the structure with the smallest distance restraint violations among the 25 generated was then selected and minimized with the CHARMM 19 SASA implicit solvation force field.^45^ All peptides where then ranked according to their binding energy, ∆*E* = *E*_*complex*_ − (*E*_*BoxA*_ + *E*_*pept*_), and the best 100 among the 40,000 generated were selected for the further analysis.

### Peptide re-docking with Glide

The ability of the 100 peptides with the best CHARMM binding energy to form complexes with the BoxA domain of HMGB1 was then additionally assessed with the peptide-docking protocol of Glide,^46^ implemented in the Schrodinger suite for molecular modeling (Version 2016-4).

Aiming to leave the algorithm free to explore the entire surface of the protein we performed, also in this case, blind docking using a grid positioned in the center of geometry (COG) and large enough to contain the entire BoxA.

For each peptide, the 15 best poses were saved for further analysis, resulting in a total of 1,500 peptide-BoxA complexes. The 200 complexes with the best Glide score were inspected and, for each peptide, only the best pose conserving some of the glycyrrhizin interactions was kept. Peptides without a glycyrrhizin-like pose in the top 200 solutions were discarded. At the end of this process, 43 peptides were discarded and 57 retained for subsequent analysis.

### Molecular dynamics (MD) and binding free energy calculations

To further asses the stability of the 57 selected complexes and to better estimate their affinity, we performed 0.5 µs MD simulations in explicit water using AMBER16. Snapshots from the corresponding trajectories were extracted to compute the binding energy ∆G with MM-GBSA, a computational method already applied in similar studies with positive results.^30, 47, 48^

All the peptide-BoxA complexes were solvated in a water box with a minimum distance from the protein surface of 10 Å. The total charge of the system was neutralized adding a proper number of Cl^−^/Na^+^ ions.

All molecular dynamics simulations were carried out using the ff14SB^49^ force field for the protein, the TIP3P model^50^ for water, and the parameters proposed by Joung et al.^51^ for the counter-ions. The peptide-BoxA complexes were first relaxed with a two-step computational protocol consisting of an energy minimization for 10,000 steps or until the energy gradient of 0.2 kcal/mol/Å^2^ was reached, restraining the backbone atomic coordinates with a harmonic restraint (k = 20 kcal/mol/Å^2^), followed by an unrestrained energy minimization for 100,000 steps (or until an energy gradient of 0.0001 kcal/mol/Å^2^ was reached). The systems were then heated to their final temperature of 300K in 40 ps. All simulations were run at constant volume, restraining the backbone coordinates (k = 20 kcal/mol/Å^2^) during the first 20 ps. Subsequently, the velocities were assigned again, and the systems equilibrated for 20ps at constant pressure (1 Atm). Finally, all complexes were simulated for 500 ns. All the simulations were analyzed and only those in which the peptide –BoxA complex was stable, were retained for MM-GBSA analysis. 500 snapshots selected in the more stable part of the simulation were used in the MM-GBSA calculations. Water molecules and counter-ions were stripped, while the protein and the peptide were parametrized using the same force field as in MD simulations. The polar contribution to solvation energy was computed with the Onufriev, Bashford and Case model setting the dielectric constant to 1 for the solute and 80 for the solvent.^52^ Finally, the 13 peptides (Table 1) with the best free energy ∆G were purchased and tested experimentally *in vitro*.

### Computational Alanine Scanning

The difference in affinity between the mutate peptides and HMGB1 was calculated by the residue scanning functionality of Bioluminate. Starting from the HBP08 pose obtained by docking all residues were mutated, one at the time, to alanine. The structure of the complex between the mutated HBP08 and HMGB1 was then refined by the side-chain prediction and backbone minimization procedure. Finally, the change in the binding free energy (ΔΔG) has been estimated by the Prime MM-GBSA procedure (OPLS2005 force field^53^ and VSGB2.1 solvent model).^31^

### Proteins and peptides

CXCL12 was chemically synthesized as previously.^54^ Histidine tagged HMGB1 was produced at the Institute of Research in Biomedicine Protein Facility (Bellinzona, Switzerland) as previously described,^12^ and stored in phosphate-buffered saline (PBS; D8537, Sigma Aldrich, Saint Louis, MO, USA). All the peptides were custom-synthesized and HPLC-purified by GenScript (New Jersey, USA). Peptides were reconstituted with DMSO and stored at −20 ºC. HPLC-MS was used to confirm 98% or higher purity for each peptide.

### Cells

A murine 300.19 PreB cell line stably transfected with the human CXCR4 was kept in culture in RPMI-1640, supplemented with 10% Fetal Bovine Serum, 1x non-essential amino acids, 1 mM Sodium pyruvate, 20 mM GlutaMAX, 50 µM β-Mercaptoethanol, 50 U/ml Penicillin and 50 µg/ml Streptomycin (GIBCO). Human monocytes were freshly isolated from buffy-coats obtained from spontaneous donation from healthy individuals (Schweizerisches Rotes Kreuz, Basel), using positive selection with CD14 microbeads (Miltenyi Biotec), as previously described.^7^

### Chemotaxis assay

Chemotaxis was performed using Boyden chambers with 5µm pore membranes, as previously described ^55^. Murine 300.19 PreB cells stably transfected with the human CXCR4, or freshly isolated human monocytes were allowed to migrate for 90 min at 37°C in response to a sub-optimal CXCL12 concentration (1 nM), in the presence or absence of HMGB1 (300 nM), as previously described ^7^. Inhibition of the synergistic activity of the CXCL12/HMGB1 heterocomplex was obtained by incubating CXCL12 and HMGB1 with 200 µM glycyrrhizin (Sigma Aldrich), as positive control^7^. All peptides, at 100 µM, were incubated with CXCL12 and HMGB1 before assessing chemotaxis, to evaluate their ability to interfere with the heterocomplex formation and inhibit the synergistic effect of HMGB1.

### Assessment of peptides toxicity

Peptides toxicity was assessed on the murine 300.19 PreB cell line expressing the human CXCR4, and on human monocytes. Cells were incubated for 2h in the presence of the different peptides at 100 µM, stained by AnnexinVFITC/Propidium Iodide following manufacturer’s instructions, and cell viability analyzed by flow cytometry in comparison to the untreated control.

### Cytokines quantification

Human monocytes were incubated for 8h at 37°C at a density of 1×10^6^ cell/ml in RPMI-1640 supplemented with 0.05% pasteurized human albumin in the presence or absence of 20nM HMGB1. A polyclonal neutralizing antibody against TLR4 (AF1478, R&D System,) was used to block TLR4 engagement. HBP08 at 100 µM was tested for its ability to inhibit HMGB1/TLR4-mediated release of cytokines. Quantification of IL1β, IL6, IL8, IL10, IL12, and TNF in the supernatants was determined by using Cytometric Bead Array (CBA) - Human Inflammatory Cytokines Kit (551811, BD Biosciences, San Jose, CA, USA), that allows the determination of the indicated human cytokines simultaneously. Acquisition was performed with FACSCanto II (BD Biosciences, San Jose, CA), and the concentration was calculated from the MFI according to a standard curve of each cytokine.

### Affinity determination by Microscale thermophoresis (MST)

The binding affinity (K_d_) between HMGB1 and the HBP08 peptide was measured by microscale thermophoresis (MST).^56^

Briefly, histidine tagged HMGB1 was labeled by the his-tag specific NT-647 dye (Monolith NTTM Protein Labelling Kit RED-NHS, NanoTemper® Technologies GmbH, Mu□nchen, Germany), for 30 minutes at room temperature. A fixed concentration of labeled HMGB1 (20nM) was mixed with 16 1:1 serial dilution of the HBP08 peptide (range 5mM-0.15 nM). The protein and the peptide were incubated for 15 minutes at room temperature, to allow binding. MST analysis was performed using premium-coated capillary tubes on a NanoTemper instrument using the following experimental settings: LED power of 5% (for fluorescence excitation), and laser power 40% (to create temperature gradient). *K*_d_ values were calculated from compound concentration-dependent changes in normalized fluorescence (Fnorm).

At least two independent experiments were performed to calculate the K_d_ values. Data were analyzed by the NanoTemper analysis software.

### Statistical analysis

The statistical significance between more than two groups was calculated by using one-way ANOVA followed by Dunnett’s multicomparisons test or two-way ANOVA followed by Tukey’s multicomparisons test.

## Acknowledgments

The authors thank Dr. Laurent Perez from the IRB protein facility for the synthesis of HMGB1. This study was supported by the Swiss National Science Foundation (3100A0-143718/1 to M.U. and 31003A-166472 to A.C.). M.U. has received funding for this project from the European Union’s Programs for research, technological development and demonstration under grant agreements ADITEC – 280873 (FP7), and TIMER – 281608 (FP7). Further supports were given by the San Salvatore Foundation, the Novartis Foundation, the Gottfried and Julia Bangerter-Rhyner-Foundation, and the Helmut Horten Foundation. A.C. has received founding for this project Swiss Cancer League (KLS-3839-02-2016-R).

## Author contributions

J.S. designed the computational pipeline, performed and analyzed the simulations and MST experiments, wrote the manuscript. V.C. designed the *in vitro* experiments, performed chemotaxis, assessed the toxicity of the peptides and their activity on TLR4, wrote the manuscript. E.M.A.F. performed and analyzed the simulations. G.D.A. performed chemotaxis experiments and assessed the toxicity of the peptides. M.G. performed MST experiments. G.D. performed chemotaxis experiments. G.G. contributed to computational design and MST experiments. M.U. designed the experiments and supervised the work, wrote the manuscript. A.C. designed the computational pipeline and supervised the work, performed simulations, wrote the code for initial peptide docking, wrote the manuscript. All the authors discussed and reviewed the manuscript.

## References

1. Charo IF, Ransohoff RM. The many roles of chemokines and chemokine receptors in inflammation. N. Engl. J. Med. 354, 610–621 (2006).

2. Muller CE, Schiedel AC, Baqi Y. Allosteric modulators of rhodopsin-like G protein-coupled receptors: opportunities in drug development. Pharmacol. Ther. 135, 292–315 (2012).

3. Wells TN, Power CA, Shaw JP, Proudfoot AE. Chemokine blockers--therapeutics in the making? Trends Pharmacol. Sci. 27, 41–47 (2006).

4. Griffith JW, Sokol CL, Luster AD. Chemokines and chemokine receptors: positioning cells for host defense and immunity. Annu. Rev. Immunol. 32, 659–702 (2014).

5. Cecchinato V, D’Agostino G, Raeli L, Uguccioni M. Chemokine interaction with synergy-inducing molecules: fine tuning modulation of cell trafficking. J. Leukoc. Biol. 99, 851–855 (2016).

6. Proudfoot AE, Uguccioni M. Modulation of Chemokine Responses: Synergy and Cooperativity. Front. Immunol. 7, 183 (2016).

7. Schiraldi M, et al. HMGB1 promotes recruitment of inflammatory cells to damaged tissues by forming a complex with CXCL12 and signaling via CXCR4. J. Exp. Med. 209, 551–563 (2012).

8. Cecchinato V, et al. Redox-Mediated Mechanisms Fuel Monocyte Responses to CXCL12/HMGB1 in Active Rheumatoid Arthritis. Front. Immunol. 9, (2018).

9. Lotze MT, Tracey KJ. High-mobility group box 1 protein (HMGB1): nuclear weapon in the immune arsenal. Nat. Rev. Immunol. 5, 331–342 (2005).

10. Andersson U, Tracey KJ. HMGB1 is a therapeutic target for sterile inflammation and infection. Annu. Rev. Immunol. 29, 139–162 (2011).

11. Yang H, Antoine DJ, Andersson U, Tracey KJ. The many faces of HMGB1: molecular structure-functional activity in inflammation, apoptosis, and chemotaxis. J. Leukoc. Biol. 93, 865–873 (2013).

12. Venereau E, et al. Mutually exclusive redox forms of HMGB1 promote cell recruitment or proinflammatory cytokine release. J. Exp. Med. 209, 1519–1528 (2012).

13. Fassi EMA, et al. Oxidation State Dependent Conformational Changes of HMGB1 Regulate the Formation of the CXCL12/HMGB1 Heterocomplex. Comput. Struct. Biotechnol. J. 17, 886–894 (2019).

14. Vénéreau E, Ceriotti C, Bianchi ME. DAMPs from Cell Death to New Life. Front. Immunol. 6, 422–422 (2015).

15. Yang H, et al. MD-2 is required for disulfide HMGB1-dependent TLR4 signaling. J. Exp. Med. 212, 5–14 (2015).

16. Mollica L, et al. Glycyrrhizin binds to high-mobility group box 1 protein and inhibits its cytokine activities. Chem. Biol. 14, 431–441 (2007).

17. Choi HW, et al. Aspirin’s Active Metabolite Salicylic Acid Targets High Mobility Group Box 1 to Modulate Inflammatory Responses. Mol. Med. 21, 526–535 (2015).

18. De Leo F, et al. Diflunisal targets the HMGB1/CXCL12 heterocomplex and blocks immune cell recruitment. EMBO reports 0, e47788.

19. VanPatten S, Al-Abed Y. High Mobility Group Box-1 (HMGb1): Current Wisdom and Advancement as a Potential Drug Target. J. Med. Chem. 61, 5093–5107 (2018).

20. Michaeli A, et al. Computationally Designed Bispecific MD2/CD14 Binding Peptides Show TLR4 Agonist Activity. J. Immunol. 201, 3383–3391 (2018).

21. Petta I, Lievens S, Libert C, Tavernier J, De Bosscher K. Modulation of Protein-Protein Interactions for the Development of Novel Therapeutics. Mol. Ther. 24, 707–718 (2016).

22. Bruzzoni-Giovanelli H, Alezra V, Wolff N, Dong CZ, Tuffery P, Rebollo A. Interfering peptides targeting protein-protein interactions: the next generation of drugs. Drug Discov. Today 23, 272–285 (2018).

23. Rader AFB, et al. Orally Active Peptides: Is There a Magic Bullet? Angew. Chem. Int. Ed. 57, 14414–14438 (2018).

24. Lee AC, Harris JL, Khanna KK, Hong JH. A Comprehensive Review on Current Advances in Peptide Drug Development and Design. Int. J. Mol. Sci. 20, (2019).

25. Lau JL, Dunn MK. Therapeutic peptides: Historical perspectives, current development trends, and future directions. Biorg. Med. Chem. 26, 2700–2707 (2018).

26. Tubert-Brohman I, Sherman W, Repasky M, Beuming T. Improved docking of polypeptides with Glide. J. Chem. Inf. Model. 53, 1689–1699 (2013).

27. Rastelli G, Del `Rio A, Degliesposti G, Sgobba M. Fast and accurate predictions of binding free energies using MM-PBSA and MM-GBSA. J. Comput. Chem. 31, 797–810 (2010).

28. Grazioso G, et al. Design of novel alpha7-subtype-preferring nicotinic acetylcholine receptor agonists: application of docking and MM-PBSA computational approaches, synthetic and pharmacological studies. Bioorg. Med. Chem. Lett. 19, 6353–6357 (2009).

29. Geng L, et al. Structure-based Design of Peptides with High Affinity and Specificity to HER2 Positive Tumors. Theranostics 5, 1154–1165 (2015).

30. Lammi C, Sgrignani J, Roda G, Arnoldi A, Grazioso G. Inhibition of PCSK9(D374Y)/LDLR Protein-Protein Interaction by Computationally Designed T9 Lupin Peptide. ACS Med. Chem. Lett. 10, 425–430 (2019).

31. Beard H, Cholleti A, Pearlman D, Sherman W, Loving KA. Applying Physics-Based Scoring to Calculate Free Energies of Binding for Single Amino Acid Mutations in Protein-Protein Complexes. PloS one 8, e82849 (2013).

32. Schrödinger Release 2018-4: BioLuminate, Schrödinger, LLC, New York, NY, 2018.

33. Arranz-Gibert P, Ciudad S, Seco J, García J, Giralt E, Teixidó M. Immunosilencing peptides by stereochemical inversion and sequence reversal: retro-D-peptides. Sci. Rep. 8, 6446 (2018).

34. Liu M, et al. D-Peptides as Recognition Molecules and Therapeutic Agents. Chem. Rec. 16, 1772–1786 (2016).

35. Pitzalis C, Kelly S, Humby F. New learnings on the pathophysiology of RA from synovial biopsies. Curr. Opin. Rheumatol. 25, 334–344 (2013).

36. Smolen JS, et al. Rheumatoid arthritis. Nat. Rev. Dis. Primers 4, 18001 (2018).

37. Fosgerau K, Hoffmann T. Peptide therapeutics: current status and future directions. Drug Discov. Today 20, 122–128 (2015).

38. He M, Bianchi ME, Coleman TR, Tracey KJ, Al-Abed Y. Exploring the biological functional mechanism of the HMGB1/TLR4/MD-2 complex by surface plasmon resonance. Mol. Med. 24, 21–21 (2018).

39. Qin S, et al. Role of HMGB1 in apoptosis-mediated sepsis lethality. J. Exp. Med. 203, 1637–1642 (2006).

40. Friesner RA, et al. Glide: a new approach for rapid, accurate docking and scoring. 1. Method and assessment of docking accuracy. J. Med/ Chem. 47, 1739–1749 (2004).

41. Stein EG, Rice LM, Brunger AT. Torsion-angle molecular dynamics as a new efficient tool for NMR structure calculation. J. Magn. Reson. 124, 154–164 (1997).

42. Guntert P, Mumenthaler C, Wuthrich K. Torsion angle dynamics for NMR structure calculation with the new program DYANA. J. Mol. Biol. 273, 283–298 (1997).

43. Fu B, et al. ALMOST: an all atom molecular simulation toolkit for protein structure determination. J. Comput. Chem. 35, 1101–1105 (2014).

44. Dominguez C, Boelens R, Bonvin AM. HADDOCK: a protein-protein docking approach based on biochemical or biophysical information. J. Am. Chem. Soc. 125, 1731–1737 (2003).

45. Ferrara P, Apostolakis J, Caflisch A. Evaluation of a fast implicit solvent model for molecular dynamics simulations. Proteins 46, 24–33 (2002).

46. Celie PHN, et al. Crystal structure of acetylcholine-binding protein from Bulinus truncatus reveals the conserved structural scaffold and sites of variation in nicotinic acetylcholine receptors. J. Biol. Chem. 280, 26457–26466 (2005).

47. Lammi C, Zanoni C, Aiello G, Arnoldi A, Grazioso G. Lupin Peptides Modulate the Protein-Protein Interaction of PCSK9 with the Low Density Lipoprotein Receptor in HepG2 Cells. Sci. Rep. 6, 29931 (2016).

48. Ylilauri M, Pentikainen OT. MMGBSA as a tool to understand the binding affinities of filamin-peptide interactions. J. Chem. Inf. Mode. 53, 2626–2633 (2013).

49. Maier JA, Martinez C, Kasavajhala K, Wickstrom L, Hauser KE, Simmerling C. ff14SB: Improving the Accuracy of Protein Side Chain and Backbone Parameters from ff99SB. J. Chem. Inf. Model. 11, 3696–3713 (2015).

50. Jorgensen WL, Chandrasekhar J, Madura JD, Impey RW, Klein LM. Comparison of simple potential functions for simulating liquid water. J. Chem. Phys. 79, 926–935 (1983).

51. Joung IS, Cheatham TE. Determination of alkali and halide monovalent ion parameters for use in explicitly solvated biomolecular simulations. J. Phys. Chem. B. 112, 9020–9041 (2008).

52. Onufriev A, Bashford D, Case DA. Exploring protein native states and large-scale conformational changes with a modified generalized born model. Proteins 55, 383–394 (2004).

53. Banks JL, et al. Integrated Modeling Program, Applied Chemical Theory (IMPACT). J. Comput. Chem. 26, 1752–1780 (2005).

54. Clark-Lewis I, Vo L, Owen P, Anderson J. Chemical synthesis, purification, and folding of C-X-C and C-C chemokines. Methods in enzymology 287, 233–250 (1997).

55. Uguccioni M, D’Apuzzo M, Loetscher M, Dewald B, Baggiolini M. Actions of the chemotactic cytokines MCP-1, MCP-2, MCP-3, RANTES, MIP-1 alpha and MIP-1 beta on human monocytes. Eur. J. Immunol. 25, 64–68 (1995).

56. Jerabek-Willemsen M, Wienken CJ, Braun D, Baaske P, Duhr S. Molecular interaction studies using microscale thermophoresis. Assay Drug Dev. Technol. 9, 342–353 (2011).

