## Supplementary Materials for "Systematic development of peptide inhibitors targeting the CXCL12/HMGB1 interaction" (from another version of this preprint: version 2, 2020-01-13)

**Supplementary information**

**Table S1.** Results of the affinity prediction, performed by MM-GBSA, for the 57 peptides selected after docking calculations. The unbound states indicate that the peptide did not conserve its original binding mode over the simulation time.

| **N°** | **Sequence** | **Simulation** | **∆G_GB_** | **Std.Error** | **Code** |
| --- | --- | --- | --- | --- | --- |
| 01 | HEMYWEDEW | OK | -52.8 | 0.3 | HBP01 |
| 02 | IDLRFFMRQ | OK | -52.0 | 0.3 | HBP02 |
| 03 | FAFELIQTD | OK | -51.7 | 0.4 | HBP03 |
| 04 | CIPMMMHAW | OK | -50.0 | 0.3 | HBP04 |
| 05 | WISNWILMW | OK | -45.9 | 0.3 | HBP05 |
| 06 | TWNIHFADH | OK | -45.6 | 0.5 | HBP06 |
| 07 | HWTLANWCR | OK | -45.2 | 0.4 | HBP07 |
| 08 | GYHYERWIH | OK | -45.1 | 0.5 | HBP08 |
| 09 | QFMKNCEEM | OK | -44.8 | 0.4 | HBP09 |
| 10 | SINWHMYVN | OK | -44.8 | 0.3 | HBP10 |
| 11 | MYRENQPTR | OK | -42.9 | 0.4 | HBP11 |
| 12 | YHICWYGDY | OK | -42.5 | 0.4 | HBP12 |
| 13 | WLWYEWGWQ | OK | -41.9 | 0.3 | HBP13 |
| 14 | DYCWKIMTQ | OK | -41.9 | 0.3 |  |
| 15 | WCHFFFPHW | OK | -41.6 | 0.4 |  |
| 16 | MKSSDCCLE | OK | -39.7 | 0.5 |  |
| 17 | EWFVMKHLN | OK | -39.0 | 0.4 |  |
| 18 | MIRDQILHN | OK | -38.9 | 0.4 |  |
| 19 | WHQLTEHWI | OK | -38.2 | 0.5 |  |
| 20 | HDHDFWAWY | OK | -37.5 | 0.2 |  |
| 21 | WQWHQFQGR | OK | -35.9 | 0.3 |  |
| 22 | VMASWQHGL | OK | -34.9 | 0.5 |  |
| 23 | LDNFLGDHW | OK | -34.7 | 0.4 |  |
| 24 | PRMGWEKPE | OK | -34.1 | 0.4 |  |
| 25 | WICVWHHAS | OK | -33.9 | 0.3 |  |
| 26 | IRWCVDARY | OK | -30.0 | 0.6 |  |
| 27 | WNAMSFCCS | OK | -28.9 | 0.4 |  |
| 28 | IFHIMTEMW | OK | -28.7 | 0.2 |  |
| 29 | FDRPRYRTT | OK | -28.6 | 0.4 |  |
| 30 | QIEDMPTSK | OK | -28.3 | 0.4 |  |
| 31 | FDCMMDMTK | OK | -28.0 | 0.3 |  |
| 32 | NTVALKLRD | OK | -27.8 | 0.4 |  |
| 33 | YHYHMLMQS | OK | -27.7 | 0.3 |  |
| 34 | NITHNVWHR | OK | -27.6 | 0.3 |  |
| 35 | DRNLEVEQI | OK | -26.7 | 0.3 |  |
| 36 | HYNKWKHQE | OK | -25.6 | 0.4 |  |
| 37 | ICMPPNTKN | OK | -24.6 | 0.3 |  |
| 38 | SMIPVQEAS | OK | -24.5 | 0.3 |  |
| 39 | YQRNELEYL | OK | -24.4 | 0.2 |  |
| 40 | HYFDMLHFH | OK | -21.0 | 0.4 |  |
| 41 | SHYFKHSNF | OK | -19.8 | 0.4 |  |
| 42 | FIKQMEEST | OK | -18.5 | 0.3 |  |
| 43 | KYQWMHYTP | OK | -16.9 | 0.6 |  |
| 44 | FVGMRWKFL | Unbound | / | / |  |
| 45 | WQIPDHRDH | Unbound | / | / |  |
| 46 | QCFHPSFED | Unbound | / | / |  |
| 47 | VPSSAKNRD | Unbound | / | / |  |
| 48 | KHMTKCEQW | Unbound | / | / |  |
| 49 | ETYQFRPNK | Unbound | / | / |  |
| 50 | WNCHRDRPK | Unbound | / | / |  |
| 51 | KHMTKCEQW | Unbound | / | / |  |
| 52 | KCVVFHYDP | Unbound | / | / |  |
| 53 | PTFEEFAAF | Unbound | / | / |  |
| 54 | QCFHPSFED | Unbound | / | / |  |
| 55 | EWLYRQEYH | Unbound | / | / |  |
| 56 | QDYAPRASN | Unbound | / | / |  |
| 57 | KDKAFKNVS | Unbound | / | / |  |

Std.Error = Standard Error of Mean

**Figure S1.** Representation of docking structures between Box-A of HMGB1 (cyan) and tested peptides (different colors) from HBP01 (A) to HBP13 (M).

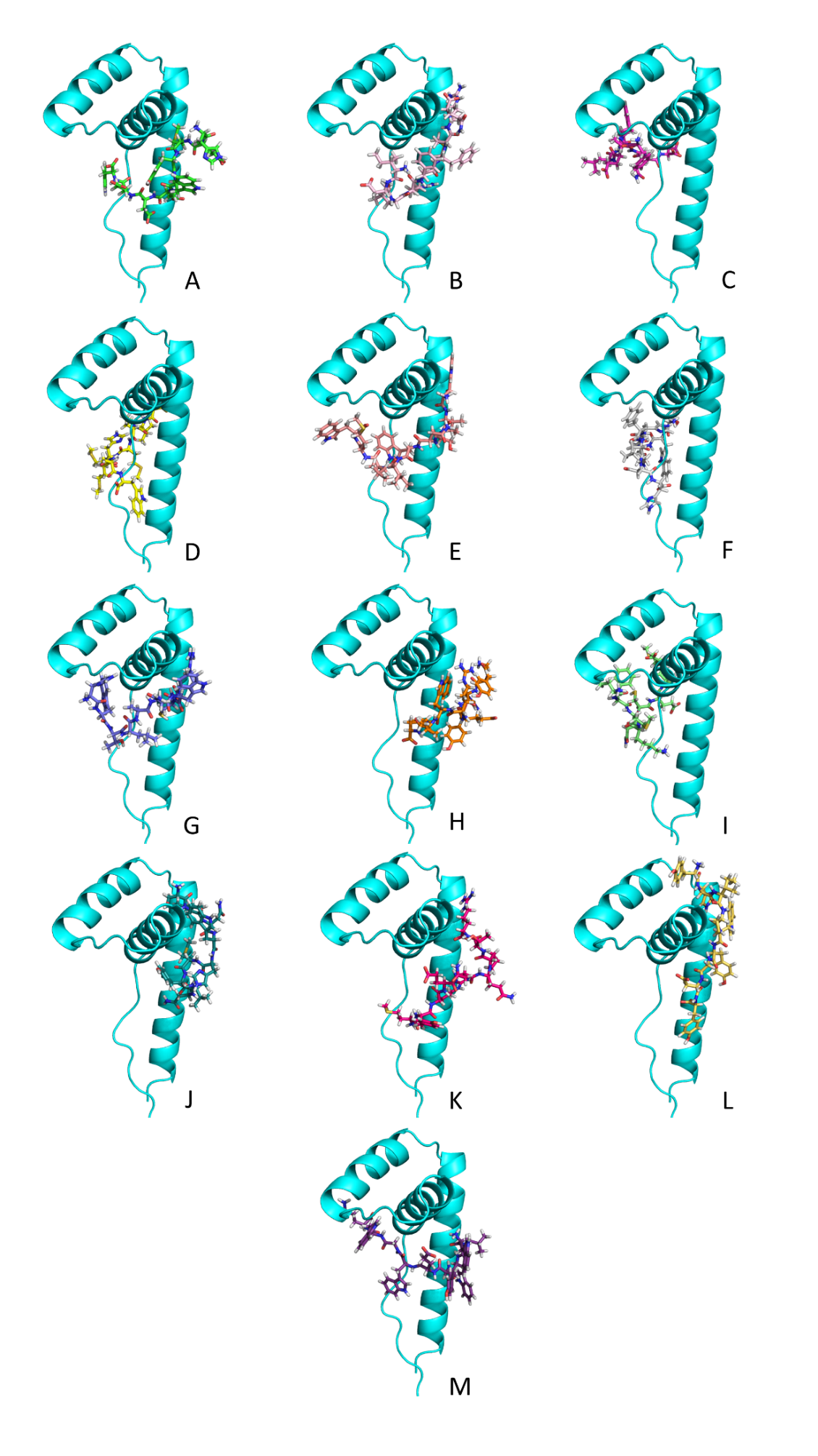
